# Dissociating the intensity and phase origins of sleepiness through a threshold-distance model of sleep-wake dynamics

**DOI:** 10.64898/2026.08.10.743860

**Authors:** Yi Yao, Zijun Ning, Dongping Yang, Chenggui Yao

**Affiliations:** College of Data Science, Jiaxing University, Jiaxing 314000, China iversity of Chinese Academy of Sciences, Wenzhou 325001, China; Shenzhen Loop Area Institute (SLAI), Shenzhen 518045, China

## Abstract

Sleepiness is a leading proximate cause of drowsy-driving fatalities, medical errors and industrial accidents, yet it has resisted mechanistic prediction; although it arises from well-characterized sleep-wake physiology, it is experienced as a subjective state and has lacked a quantitative link to the underlying dynamics. We previously showed that subjective sleepiness maps linearly, with a protocol-invariant form, onto the signed distance *H − H*^+^ between the homeostatic pressure *H* and the circadian-modulated sleep-onset threshold *H*^+^. This single quantity predicts sleepiness accurately but is mechanistically ambiguous: the same value can arise either because *H* sits far from the boundary or because the threshold *H*^+^(*t*) has shifted with circadian phase, and these two origins call for entirely different interpretations and interventions. Here we resolve this ambiguity by decomposing *H − H*^+^ into two mechanistically separable axes–*intensity* and *phase*. The *intensity* axis is the time-averaged margin ⟨ *H − H*^+^⟩, set by how far, on average, *H* sits from the sleep boundary: slowed homeostatic accumulation accounts for the paradoxically blunted sleepiness of older adults, and pharmacological suppression of *H* accounts for the dose-dependent alerting effect of caffeine. The *phase* axis is set by the circadian modulation of *H*^+^(*t*): under a forced-desynchrony protocol, in which the pacemaker free-runs and the homeostatic and circadian processes are experimentally decoupled, sleepiness tracks the circadian profile of *H*^+^(*t*) across all phases while the intensity mapping itself remains unchanged–a clean dissociation of the two axes. By resolving felt sleepiness into these two physiological degrees of freedom, this framework renders previously isolated phenomena–aging, caffeine and circadian misalignment–commensurable within a single theory and provides a physiologically interpretable basis for prospective fatigue-risk prediction.

**Author summary:** Why people feel sleepy after sleep loss, or at particular times of day, remains difficult to predict from physiology alone. Sleep and wake are shaped by two interacting processes: a daily circadian rhythm and a homeostatic pressure that builds during wakefulness. In earlier work, we linked subjective sleepiness ratings to a simple geometric quantity–how close sleep pressure sits to a circadian sleep-onset boundary. That link is useful, but ambiguous: the same distance can arise either because pressure itself has changed, or because the boundary has moved with circadian phase. Here we use a computational model of the sleep-wake switch, extended to include the wake-stabilizing orexin system, to separate these contributions into an intensity axis and a phase axis. We find that aging and caffeine mainly alter how large the average distance to the boundary becomes, whereas forced desynchrony mainly alters how that distance varies across circadian phase. This dissociation offers a compact way to interpret several otherwise separate observations within one quantitative picture, and a step toward more physiologically grounded fatigue-risk assessment.

## Introduction

Sleepiness is among the most universal of human experiences, yet it remains one of the hardest to predict. It is the proximate cause of drowsy-driving fatalities, medical errors, and industrial accidents, and the quantity that clinical and operational fatigue-management systems ultimately seek to forecast [1]. The central difficulty is that sleepiness is a *subjective, felt* quantity, whereas the physiology that generates it–the interplay of circadian phase and accumulated sleep pressure–is objective and dynamical [2, 3]. Bridging the two has proven remarkably elusive: how does the continuous internal state of the sleep-regulatory system give rise to the discrete moment at which a person reports feeling sleepy?

The standard instrument for gauging that felt state, the Karolinska Sleepiness Scale (KSS) [1], throws this gap into sharp relief. Although the KSS is a foundational metric of subjective alertness in both clinical research and high-stakes operational environments [4], the biophysical frameworks that describe the sleep-regulatory system have never been robustly linked to it. The prevailing two-process framework and its descendants relate sleepiness to a single *sleep drive* that fuses the homeostatic (*H*) and circadian (*C*) components into one opaque number [5–7], so that the independent contributions of “how long one has been awake” and “what time it is” are irrecoverably entangled. Consequently, existing predictors–whether built on empirical linear regression or on loose associations with model-derived variables such as the total sleep drive–are effectively curve fits rather than mechanistic explanations: they offer no principled account of *why* sleepiness rises when it does, and the relationship between a biologically interpretable internal state and KSS remains open.

We argue that this difficulty dissolves once sleepiness is referred not to the drive itself, but to a threshold. In the bistable flip-flop network that governs sleep and wakefulness, the brain switches state when the homeostatic drive crosses a circadian-modulated *switching threshold*–the level at which the wake- and sleep-promoting populations abruptly exchange dominance [8, 9]. This suggests a simple and physically transparent hypothesis: subjective sleepiness reflects **how close the system currently sits to its sleep-onset threshold**. Under this view, feeling sleepy is the perceptual read-out of an impending state transition [10, 11].

Crucially, because the signed distance *H − H*^+^ is the difference of two physiologically distinct variables–homeostatic pressure *H* and the circadian threshold *H*^+^–the same geometric quantity that predicts sleepiness also *dissociates* it into two axes that conventional fused-drive accounts cannot separate: an *intensity* axis, set by how far *H* sits from the boundary, and a *phase* axis, set by when *H*^+^(*t*) rises and falls.

That the switching threshold is not merely a theoretical construct but an operationally consequential quantity is underscored by its growing role in clinical and personalized sleep medicine. By defining the position of an individual’s homeostatic drive relative to the state-switching thresholds, Hong et al. constructed a sleep-sufficiency metric that quantifies how adequately shift workers recover from accumulated sleep debt, and used it to design personalized schedules that alleviate daytime sleepiness [12]; Building on the same threshold logic, Song et al. delivered real-time, model-guided sleep interventions through wearable devices [13]; Skeldon et al. leveraged threshold-referenced longitudinal data to disambiguate whether an individual’s sleep phenotype is governed by endogenous circadian rhythms or environmental light exposure [7]. Furthermore, the Homeostatic–Circadian–Light model further demonstrated that explicitly modeling both sleep and wake thresholds substantially improves predictions of sleep timing and duration across varying environmental light conditions [14].

Across these diverse applications the thresholds serve a common function: they constitute the critical interface between physiology and behavior, and shifting them alters the phase, duration, and consolidation of sleep. Yet in every case the thresholds employed are either phenomenological parameters inherited from the two-process framework or fixed quantities within prior flip-flop formulations–none incorporates the orexinergic drive that is the principal physiological determinant of threshold position and state stability [15, 16]. This disconnect means that current personalized models can adjust for circadian phase and homeostatic load but cannot account for the inter-individual and age-related variation in state-holding capacity that originates in the orexin system.

Building on our previous demonstration that subjective sleepiness maps linearly onto the geometric distance *H − H*^+^ [17], the present work has two aims. First, we recertify that protocol-invariant linear law in a physiologically extended sleep–wake network that includes explicit orexinergic excitation of the wake-promoting population [8, 15], so that *H*^+^ is a derived, rather than a fitted, boundary, and verify that KSS = *a* (*H − H*^+^) + *b* continues to hold under both acute total deprivation and chronic sleep restriction. Second–and this is the central conceptual advance–we show that the same distance resolves into two separable control axes: **intensity**–how large ⟨ *H − H*^+^ ⟩ becomes–is set primarily by homeostatic accumulation of *H* (and by steady shifts of the threshold that change the mean margin), whereas **phase**–when sleepiness peaks and troughs within the day–is set by the circadian modulation of *H*^+^(*t*).

Within this framework we address four problems that conventional accounts treat separately. First, we recertify the linear law against behavioral data with the orexin-dependent sleep-onset threshold—the quantitative backbone of all subsequent claims. Second, we dissect the *intensity* axis in aging: older adults’ blunted post-deprivation sleepiness is reproduced by slowed homeostatic accumulation, with circadian amplitude acting instead on the temporal patterning of KSS rather than its mean level. Third, we show that caffeine acts on the same intensity axis by pharmacologically suppressing effective *H*, shifting the operating point down the identical linear curve without changing its slope [18]. Finally, we isolate the *phase* axis under forced desynchrony: by scheduling sleep and wake to a non-24 h day, the pacemaker is allowed to free-run so that *H* becomes decoupled from the circadian phase of *H*^+^(*t*); sleepiness then tracks the circadian modulation of the threshold across all phases, exposing the phase contribution of *C* while the intensity mapping is held fixed [19]. In each case, the threshold distance supplies a common quantitative language for outcomes that are otherwise modeled in isolation, transforming a heterogeneous collection of empirical observations into predictions of a single, mechanistically grounded theory.

## Results

### An extended sleep–wake model and the threshold distance as a metric of subjective sleepiness

To evaluate the threshold-distance hypothesis quantitatively, we require a dynamical substrate in which both the homeostatic pressure *H* and the sleep-onset threshold *H*^+^ are physiologically determined rather than fitted by hand. We therefore adopt the mean-field sleep-wake network established [6, 8, 20], which extends the Phillips-Robinson framework by resolving orexin (Orx) as an explicit wake-promoting population rather than subsuming it into a lumped monoaminergic system. The model comprises three interacting neural populations–the sleep-promoting ventrolateral preoptic area (VLPO), an aggregate non-Orx monoaminergic wake-promoting population (MA), and Orx–whose connectivity is shown schematically in Fig. 1a. Sleep-wake switching is regulated by a circadian drive *C* (SCN-derived rhythmic input entrained by the light-dark cycle) and a homeostatic drive *H* (sleep pressure accumulating during wakefulness and dissipating during sleep, in accord with the two-process framework [2]). The dynamical equations for the full sleep model (Fig. 1b) and parameter values (Tab. 1) are provided in Methods; those parameters are set to yield a stable baseline of regular sleep-wake cycling with sharp state transitions. We use this substrate only insofar as it yields well-defined trajectories of *H*(*t*) and *H*^+^(*t*): the scientific claims below concern the geometric quantity *H− H*^+^ and its intensity/phase dissociation (Figs. 2a–b).

**Fig 1.**
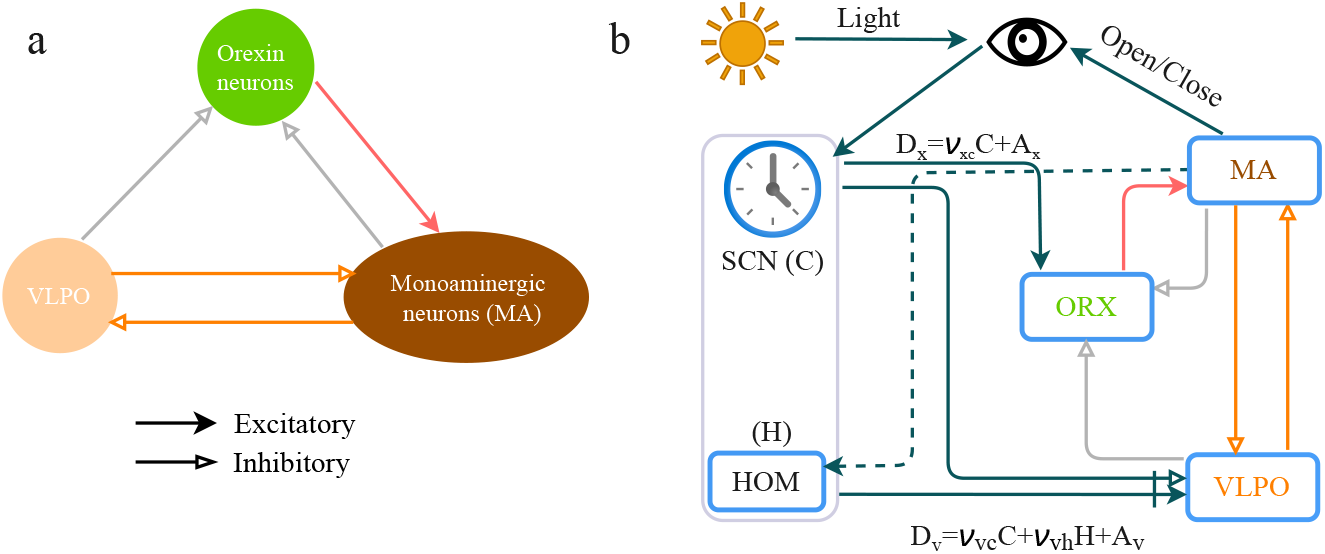
Schematic of the extended Phillips–Robinson sleep–wake model [20]. (a) Three mutually interacting neural populations: the sleep-active ventrolateral preoptic area (VLPO), an aggregate monoaminergic wake-promoting population (MA), and the orexinergic neurons (Orx) of the lateral hypothalamic area [21]. Reciprocal VLPO-MA inhibition implements the canonical sleep-wake flip-flop, while Orx provides a stabilizing excitatory drive to MA and receives inhibitory feedback from VLPO. (b) The circuit is coupled to the two-process model of sleep regulation: the circadian drive *C* from the suprachiasmatic nucleus (SCN), and the homeostatic drive *H*, which accumulates during wake and dissipates during sleep, jointly modulate VLPO activity. Solid arrows denote excitatory couplings; hollow arrows denote inhibitory couplings.

**Fig 2.**
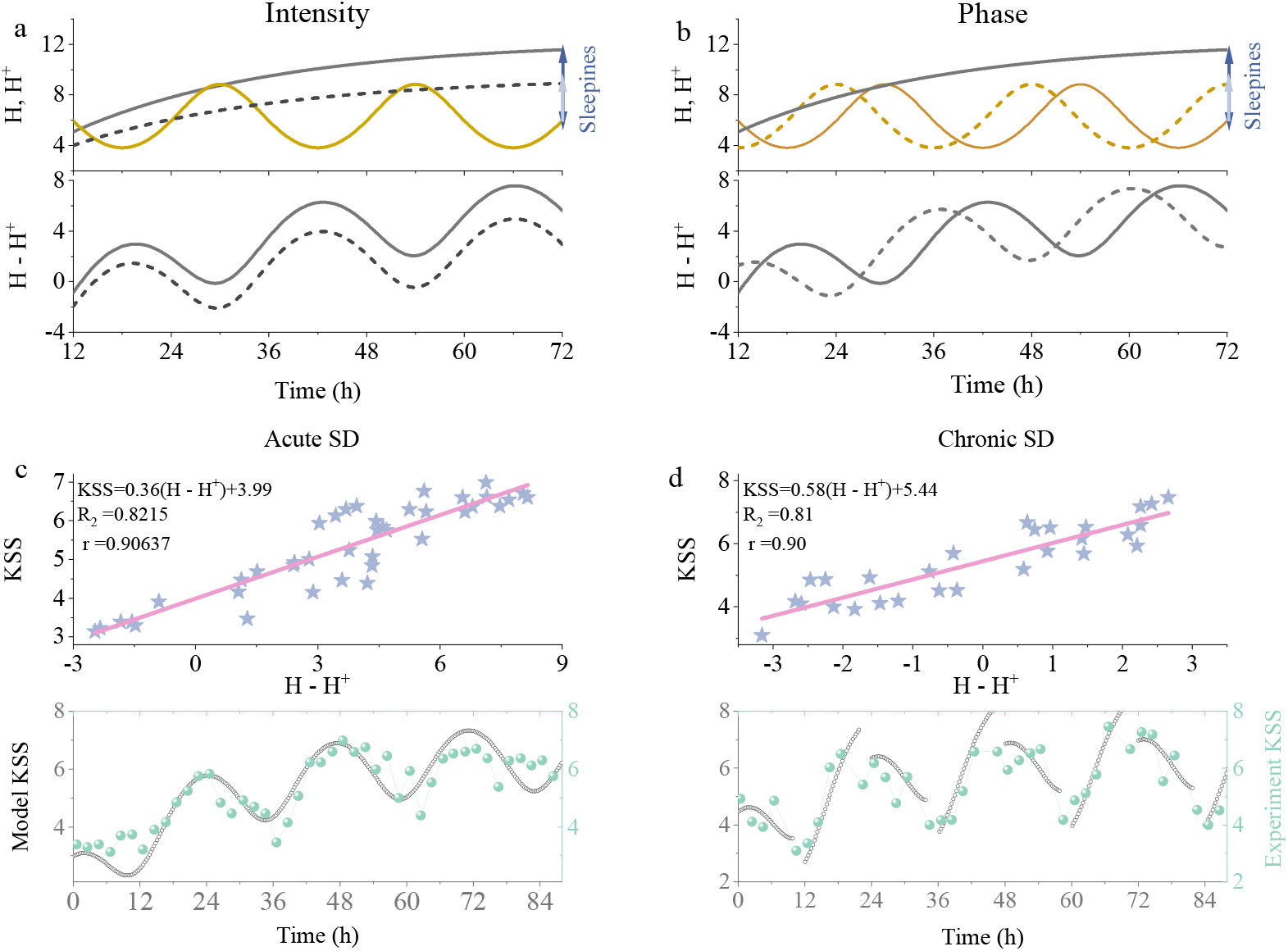
Definition and validation of the sleepiness metric. The sleepiness metric *S* = *H − H*^+^ is defined as the signed distance between the homeostatic drive *H* and the circadian sleep-onset threshold *H*^+^. (a - b) The schematic diagram for the effect of intensity level (left) and circadian phase (right) on sleepiness. (c) KSS plotted against *S* = *H − H*^+^, showing a linear relationship that holds across both protocols: the 88-h acute sleep deprivation (SD, right) and the 14-day chronic sleep restriction (2 h sleep per night, left) [22]. (d) Time courses of the measured KSS scores and the corresponding fitted *S* for acute SD (right) and chronic restriction (left).

**Table 1.**
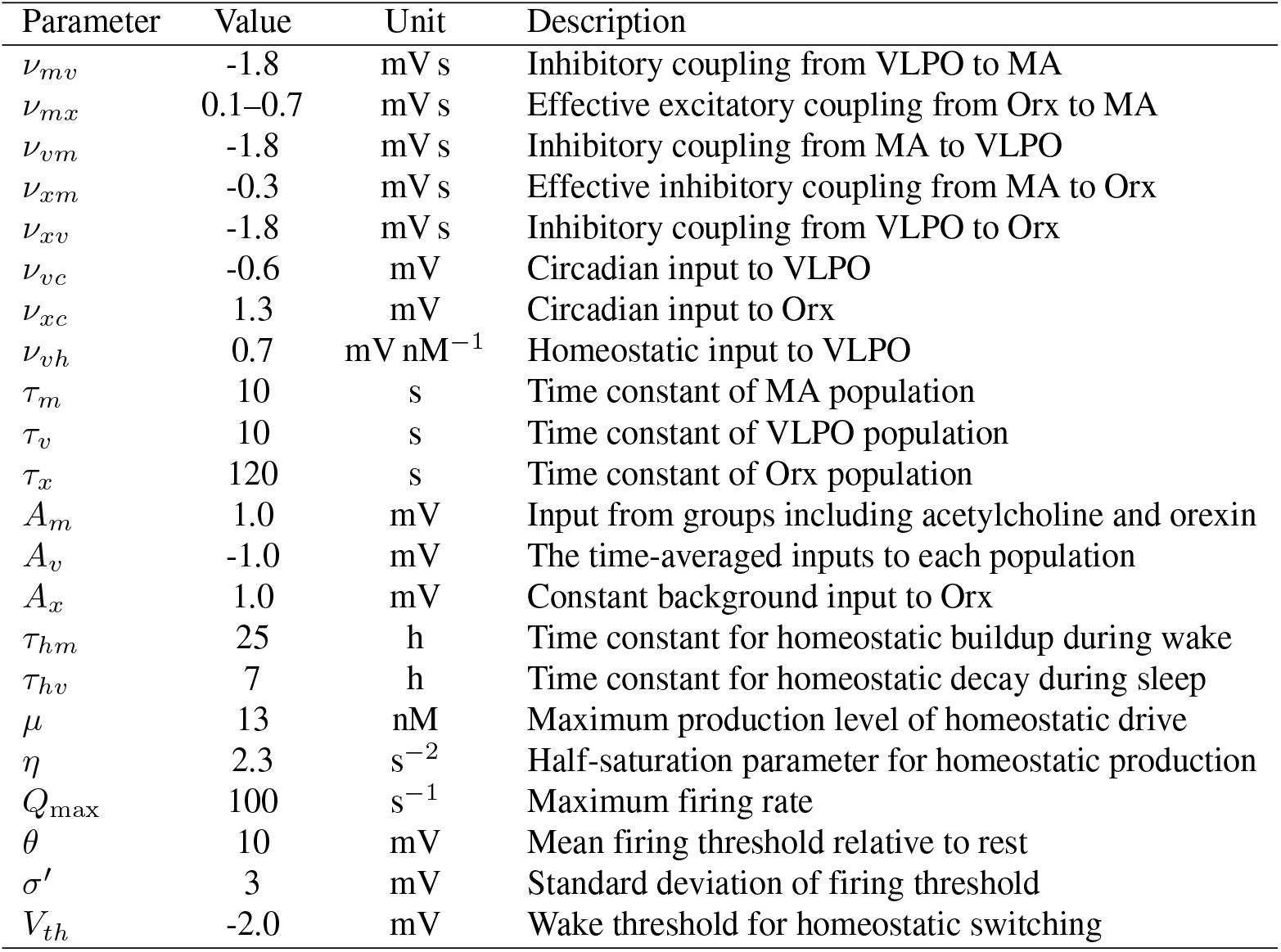
Nominal parameter values. Here, *v*_*ab*_ denotes the effective coupling from population *b* to population *a*.

| Parameter | Value | Unit | Description |
| --- | --- | --- | --- |
| $\nu_{mv}$ | -1.8 | mV s | Inhibitory coupling from VLPO to MA |
| $\nu_{mx}$ | 0.1–0.7 | mV s | Effective excitatory coupling from Orx to MA |
| $\nu_{vm}$ | -1.8 | mV s | Inhibitory coupling from MA to VLPO |
| $\nu_{xm}$ | -0.3 | mV s | Effective inhibitory coupling from MA to Orx |
| $\nu_{xv}$ | -1.8 | mV s | Inhibitory coupling from VLPO to Orx |
| $\nu_{vc}$ | -0.6 | mV | Circadian input to VLPO |
| $\nu_{xc}$ | 1.3 | mV | Circadian input to Orx |
| $\nu_{vh}$ | 0.7 | mV nM<sup>-1</sup> | Homeostatic input to VLPO |
| $\tau_m$ | 10 | s | Time constant of MA population |
| $\tau_v$ | 10 | s | Time constant of VLPO population |
| $\tau_x$ | 120 | s | Time constant of Orx population |
| $A_m$ | 1.0 | mV | Input from groups including acetylcholine and orexin |
| $A_v$ | -1.0 | mV | The time-averaged inputs to each population |
| $A_x$ | 1.0 | mV | Constant background input to Orx |
| $\tau_{hm}$ | 25 | h | Time constant for homeostatic buildup during wake |
| $\tau_{hv}$ | 7 | h | Time constant for homeostatic decay during sleep |
| $\mu$ | 13 | nM | Maximum production level of homeostatic drive |
| $\eta$ | 2.3 | s<sup>-2</sup> | Half-saturation parameter for homeostatic production |
| $Q_{\max}$ | 100 | s<sup>-1</sup> | Maximum firing rate |
| $\theta$ | 10 | mV | Mean firing threshold relative to rest |
| $\sigma'$ | 3 | mV | Standard deviation of firing threshold |
| $V_{th}$ | -2.0 | mV | Wake threshold for homeostatic switching |

In previous work, we carried out an initial exploration of how subjective sleepiness might be quantified and predicted within the reduced sleep model [17], defining sleepiness as the *signed distance* between the instantaneous homeostatic drive and the circadian sleep-onset threshold,

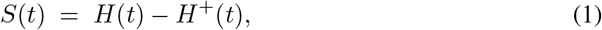

as illustrated in Fig. 2a. Moreover, we showed that the experimental KSS scores, when plotted against *H − H*^+^, collapse onto a simple linear relationship [17],

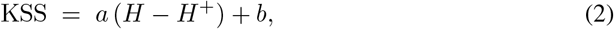

with slope *a* and intercept *b*. As detailed in Methods, imposing the wake-to-sleep boundary condition on the VLPO drive yields the new sleep-onset threshold of the extended sleep model with orexin [23],

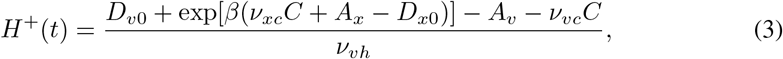

where *β, D*_*v*0_, and *D*_*x*0_ are fitting coefficients, and *C* = sin(*ωt* + *ϕ*_*c*_) denotes the circadian drive [8, 20], with the initial phase *ϕ*_*c*_ anchored to the experimental markers by requiring the model CBTmin to coincide with the measured core-body-temperature minimum *t*_CBTmin_ [5]. The meaning and values of the remaining parameters (Tab. 1) are provided in Methods. During wakefulness, the homeostatic pressure *H* evolves as

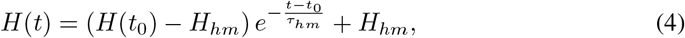

where *H*(*t*_0_) is the initial value, *H*_*hm*_ *≈* 0.94 *µ* is the saturation value and *µ* is maximum production level of homeostatic pressure, and *τ*_*hm*_ is the relaxation time constant.

To validate this linear law with the new sleep-onset threshold, we simulated two canonical sleep-deprivation protocols for young subjects: (i) an 88-hour acute sleep-deprivation (SD) protocol, in which participants underwent sustained wakefulness following three baseline days of normal sleep; and (ii) a chronic sleep-restriction protocol, in which participants underwent 14 consecutive nights of restricted sleep (2 h per night) after habituation and baseline nights, following van Dongen *et al*. [22]. Both protocols were reproduced in the model, and the resulting trajectories of the neural populations and threshold variables were computed. Sampling *H − H*^+^ at the same clock times as the behavioral measurements revealed a striking regularity: the model-derived threshold distance tracked the KSS scores with remarkable fidelity across both protocols. In each case, plotting KSS against *H− H*^+^ collapsed the time-series data onto a single linear relationship (Figs. 2(c) - 2(d)). That the same functional form–and closely similar coefficients–holds across two markedly different sleep-loss histories is nontrivial: it implies that subjective sleepiness is determined neither by cumulative sleep debt nor by elapsed wake time per se, but by the instantaneous position of the system relative to its circadian-modulated transition boundary. Different deprivation and restriction schedules thus converge onto a common low-dimensional physiological coordinate, *H− H*^+^, which governs perceived sleepiness.

### Intensity-Phase dissociation

The threshold distance therefore serves not merely as a predictive correlate but as a mechanistic state variable of the sleep–wake regulatory system. Because *H*^+^ is an explicit function of the circadian drive *C* and the neural coupling strengths (Eq. (3)), while *H* evolves according to simple first-order homeostatic dynamics (Eq. (4)), the quantity *H − H*^+^ provides a transparent decomposition of sleepiness into two operationally distinct axes that the remainder of the paper keeps separate. The *intensity* axis is the margin ⟨*H − H*^+^⟩: it asks how deeply the system sits in the sleep-promoting region and is therefore the natural target of homeostatic manipulations that move *H* (or of steady threshold shifts that change the mean boundary), as shown in Fig. 2(a). The *phase* axis is the circadian modulation of *H*^+^(*t*): it asks *when* the margin narrows and widens within the day and is therefore the natural target of entrainment and schedule interventions, as shown in Fig. 2(b). This decomposition enables analytical prediction:

⟨*H − H*^+^⟩ can be expressed in closed form as a function of physiological parameters (Eq. (19)), including the homeostatic gain, the circadian coupling strengths, and the prior wake duration. It is precisely this closed-form tractability–together with the intensity/phase split–that makes the metric portable across the caffeine, aging, and the forced-desynchrony conditions examined next.

### Case I of Intensity axis: reduced sleepiness in older adults

Having established that *H − H*^+^ faithfully encodes subjective sleepiness across distinct deprivation schedules in healthy young adults, we next asked whether the same metric can account for a well-documented but mechanistically puzzling age-related dissociation: older adults consistently report *lower* subjective sleepiness than their younger counterparts following acute sleep deprivation, despite exhibiting comparable or even greater performance impairment [24–26]. This is a probe of the *intensity* axis: what features of the sleep–wake regulatory system compress the mean margin ⟨*H − H*^+^⟩ in aging?

To address this question, we simulated a 48-h acute SD protocol in both a young-parameterized (*µ* = 13.0 nM, *τ*_*hm*_ = 25.0 h, *v*_*xc*_ = 1.3, *v*_*vc*_ = *−* 0.6) and an aging-parameterized (*µ* = 11.0 nM, *τ*_*hm*_ = 45.0 h, *v*_*xc*_ = 1.04, *v*_*vc*_ = *−* 0.48) version of the model and compared the resulting trajectories of *H* and *H*^+^ (Figs. 3a–b). The dominant age-related change is immediately apparent: homeostatic pressure accumulates substantially more slowly in the older model during sustained wakefulness, consistent with the well-characterized attenuation of sleep homeostasis in aging reported by electroencephalographic (EEG) slow-wave-activity studies [27]. This is an intensity-axis effect: it reduces how far *H* climbs above *H*^+^, and thereby compresses the excursion of *H − H*^+^ during extended wakefulness. At any given time point after sleep-deprivation onset, the threshold distance is smaller in the older model than in the young model, and consequently the predicted sleepiness is lower.

**Fig 3.**
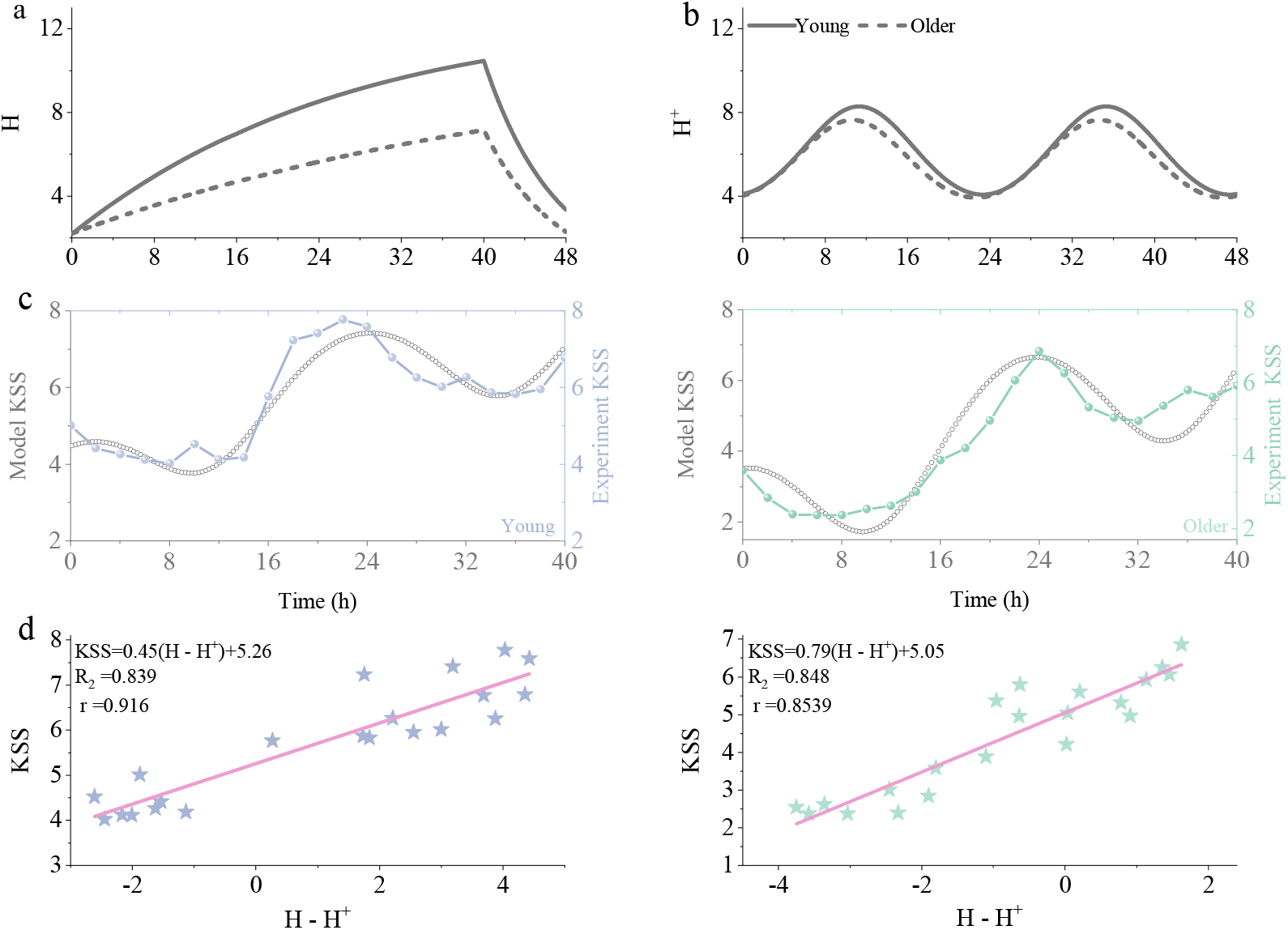
Comparison of sleepiness dynamics between young and older adults. **(a)-(b)** Time series of homeostatic pressure *H* and the sleep-onset threshold *H*^+^ for young (solid lines) and older (dashed lines) adults following 48-h sleep deprivation (SD). **(c)** Time courses of model-predicted and experimentally measured KSS scores for young (left) and older (right) adults. **(d)** KSS plotted against *H − H*^+^ for young (left) and older (right) adults, revealing a linear relationship that holds consistently across both age groups.

This age-dependent compression of *H − H*^+^ propagates directly to subjective sleepiness via the linear mapping established in Eq. (2). Figure 3c overlays the resulting model-predicted KSS time courses with the experimental KSS data [24] for both age groups. In both model and experiment, older adults maintain lower KSS scores throughout the 48-h deprivation window. Crucially, this age contrast emerges without any *ad hoc* rescaling of the KSS mapping coefficients: the same linear invariant that was validated in young adults (Fig. 2) is applied unchanged to the aging model, and the resulting predictions reproduce the experimentally observed age-related attenuation of subjective sleepiness with quantitative fidelity. Figure 3d confirms this point explicitly by plotting KSS against *H − H*^+^ for both age groups: the data collapse onto a single linear trend, demonstrating that the age-related reduction in reported sleepiness does not reflect a change in the perceptual mapping from physiological state to subjective report, but rather a genuine reduction in the magnitude of ⟨*H− H*^+^⟩. These results suggest that the lower subjective sleepiness observed in older adults after sleep loss is not paradoxical but is instead a direct and quantifiable consequence of attenuated homeostatic pressure accumulation–an intensity-axis signature of age-related changes in sleep-regulatory neurophysiology that the threshold distance *H − H*^+^ captures.

To move beyond protocol-specific comparisons and identify which physiological changes of aging most strongly govern the reduction in subjective sleepiness, we derived a closed-form expression for the time-averaged sleepiness index ⟨*H− H*^+^⟩ (Eq. (19)) and, under a fixed 40-h total-sleep-deprivation protocol, systematically varied three physiologically meaningful parameters that control the mean margin and are known to change with age.

First, reducing the circadian amplitude ratio *r*–defined as the aged-to-baseline ratio of the circadian oscillation, and thereby modeling the well-documented flattening of circadian rhythmicity in older adults [24, 27]–left the post-deprivation mean KSS essentially unchanged, producing only a marginal increase (Fig. 4a). This near-invariance shows that circadian amplitude has little leverage over the *mean intensity* of sleepiness accumulated during sustained wakefulness, and therefore cannot account for the robust age-related reduction in reported sleepiness. The attenuation of felt sleepiness in aging must instead be sought in parameters that govern homeostatic accumulation and the steady position of the sleep-onset boundary.

**Fig 4.**
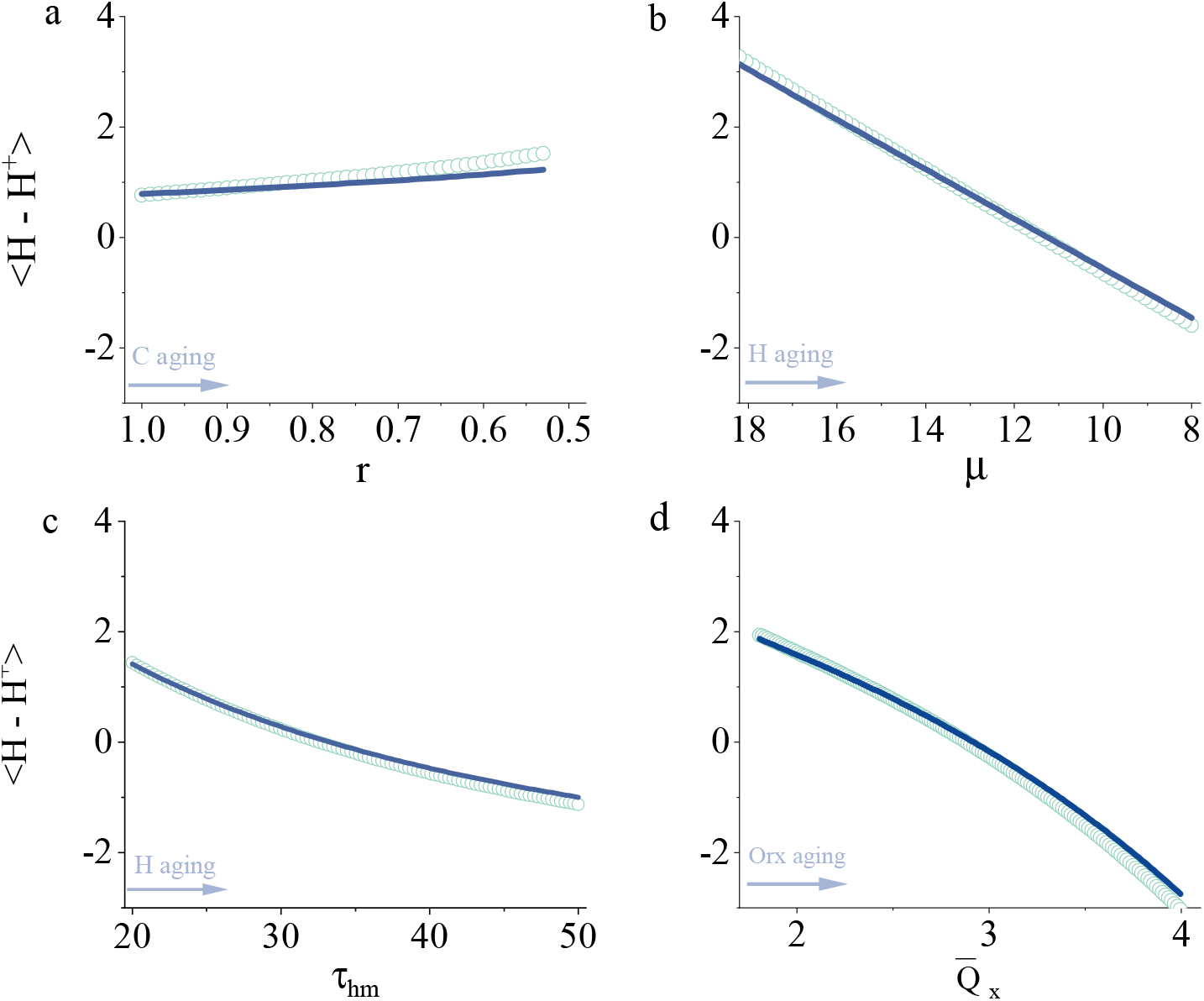
Sensitivity of model-predicted sleepiness intensity to age-related physiological parameters. Mean sleepiness after 40 h of total sleep deprivation, quantified as KSS = *a* ⟨*H − H*^+^⟩ + *b*, is shown as a function of four key parameters that change with aging: *r*, the ratio of aged to baseline circadian amplitude (phase axis: little effect on mean intensity); **(b)** *µ*, the upper asymptote of homeostatic pressure (intensity axis); **(c)** *τ*_*hm*_, the time constant governing homeostatic pressure buildup during wakefulness (intensity axis); and **(d)**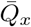, the mean firing rate of the wake-promoting orexin population (intensity axis via steady elevation of the threshold margin). Circles denote numerical simulation results; solid lines show the corresponding analytical predictions from Eq. (19). The close agreement between the two confirms that the closed-form expression captures the parametric dependence of sleepiness across the physiologically relevant aging range.

Second, decreasing the maximum attainable homeostatic pressure *µ* produced a monotonic and approximately linear reduction in ⟨*H* −*H*^+^⟩ (Fig. 4b). Because *µ* sets the asymptotic ceiling of sleep pressure that can accumulate during wakefulness, a lower ceiling limits how far *H* can encroach beyond the sleep-onset boundary *H*^+^. Given that *µ* is reduced in older individuals [28, 29], this parameter change alone predicts that younger subjects will experience markedly more intense sleepiness following deprivation than their older counterparts–consistent with the experimental observations of Duffy *et al*. [26] and Gabel *et al*. [24].

Third, lengthening the homeostatic time constant *τ*_*hm*_ likewise attenuated ⟨*H − H*^+^⟩ (Fig. 4c). A larger *τ*_*hm*_ implies that homeostatic pressure builds up more sluggishly during wakefulness; for a fixed deprivation duration, *H* therefore reaches a lower peak and remains closer to *H*^+^. Because EEG studies have consistently reported slower accumulation of slow-wave activity–a standard physiological proxy for homeostatic sleep pressure–in older adults [28, 29], an age-related lengthening of *τ*_*hm*_ is physiologically well motivated and contributes, together with the reduction in *µ*, to the blunted sleepiness response observed in aging.

Finally, we examined the role of orexinergic tone. Recent evidence indicates that, paradoxically, the orexin system becomes hyperexcitable with aging: Li *et al*. [30] demonstrated that orexin neurons in aged mice exhibit elevated spontaneous firing rates and enhanced intrinsic excitability, suggesting that the wake-promoting orexinergic drive is amplified rather than diminished in later life. To capture this effect in the model, we increased the tonic excitability parameter *A*_*x*_, which raises the average firing rate of the orexin population 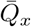. As shown in Fig. 4d, increasing 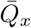 progressively lowered ⟨*H − H*^+^⟩. Mechanistically, stronger orexinergic excitation does not slow the homeostatic accumulation of *H*; rather, it raises the sleep-onset boundary *H*^+^ by reinforcing the wake-active monoaminergic drive, thereby widening the *mean* margin and reducing intensity. Age-related enhancement of orexin signalling thus provides an additional–and previously underappreciated–intensity-axis contribution to the lower subjective sleepiness observed in older adults after sleep deprivation [24, 26].

Across these parameter sweeps, the analytical predictions from Eq. (19) tracked the numerical simulations with close quantitative agreement (circles versus lines in Fig. 4a–d), validating the closed-form expression as a faithful surrogate for the full model dynamics over the physiologically relevant parameter range. The expression therefore provides a compact, transparent decomposition of ⟨*H −H*^+^⟩ into its constituent physiological determinants. Three age-sensitive factors–reduced *µ*, lengthened *τ*_*hm*_, and elevated 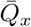 –act in the same direction on the *intensity* axis, jointly suppressing ⟨*H − H*^+^⟩, whereas circadian amplitude (*r*) exerts negligible control over mean intensity [24, 26].

### Case II of Intensity axis: reduced sleepiness by caffeine

The preceding analysis demonstrated that the linear mapping KSS = *a*(*H − H*^+^) + *b* captures age-related differences in sleepiness under caffeine-free conditions, and that those differences load primarily onto the intensity axis. A natural next step is to ask whether this threshold-based framework can also account for the well-documented alerting effect of caffeine—a second, pharmacological probe of the same intensity axis. Numerous studies have shown that coffee reduces subjective sleepiness and enhances vigilance during prolonged wakefulness [31, 32], and caffeine has been shown to attenuate EEG markers of sleep homeostasis [33, 34]. Yet, a unified quantitative description linking caffeine pharmacokinetics to the two-process sleep-wake dynamics has remained elusive. Here we extend our model to incorporate caffeine and test whether it moves sleepiness along the intensity axis without altering the phase mapping.

We reproduced the experimental design of Hansen et al. [32], in which participants maintained three baseline sleep-wake cycles before undergoing 48 h of total sleep deprivation. During the deprivation period, 200 mg caffeine was administered four times–at 13:00 and 01:30 on each of the two deprivation days–while a placebo condition served as the no-coffee control. Using the dynamical model introduced above (Eqs. (5), (12), and (14)), we first simulated the full sleep-wake history to initialize the homeostatic variable *H* at the onset of deprivation, and then evolved it forward with and without the caffeine masking term (Eq. (14)).

Figure 5(a) compares the resulting time courses of *H* for the two conditions. In the placebo case (solid lines), *H* rises monotonically toward its upper asymptote throughout the 48-h wake episode. When caffeine is present (dashed lines), each dose produces a transient but pronounced reduction in the equivalent homeostatic pressure *H*_eqv_, with the effect peaking roughly 30-60 min post-ingestion and dissipating over several hours as the drug is eliminated. The cumulative result is a markedly lower perceived sleep pressure across the deprivation period.

**Fig 5.**
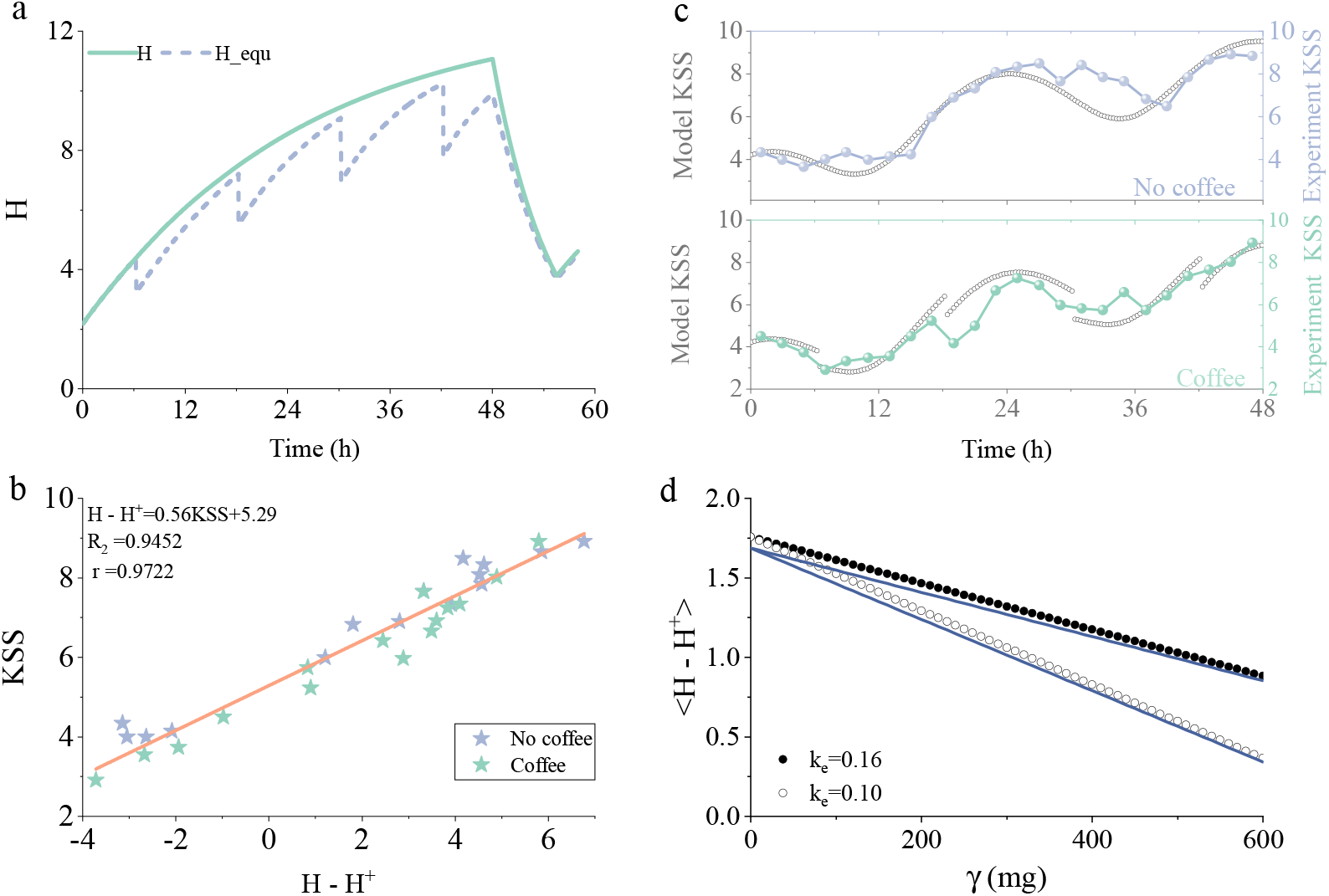
Caffeine lowers subjective sleepiness along the same threshold-distance mapping. **(a)** Time series of homeostatic pressure *H* without (solid lines) and *H*_eqv_(*t*) = *H*(*t*) [1*− ζ*_*H*_ *Z*_*C*_(*t*)] (dashed lines) with caffeine (200 mg *×* 4) during 48-h sleep deprivation (SD). **(b)** KSS versus *H− H*^+^ without (solid) and with (dashed) caffeine, collapsing onto a single linear relationship. **(c)** Time courses of model-predicted and experimentally measured KSS scores for the placebo (upper) and caffeine (lower) conditions. **(d)** Mean sleepiness after 40 h of total sleep deprivation, quantified as KSS = *a* ⟨*H − H*^+^⟩ + *b*, as a function of caffeine dose for different elimination rates *k*_*e*_.

To connect this physiological reduction to subjective experience, we sampled *H − H*^+^ at the same time points as the experimental KSS measurements. Figure 5(b) plots KSS against *H − H*^+^ for both the placebo and caffeine conditions. Remarkably, all data points–with and without coffee–collapse onto the same linear relationship established in the caffeine-free analysis. Caffeine does not alter the slope *a* or intercept *b* of the mapping; it simply shifts the operating point to lower values of *H− H*^+^ by reducing the effective homeostatic drive. This is the intensity axis in pure form: the phase structure of *H*^+^(*t*) is left intact, and only the homeostatic coordinate moves. Taken together with the aging results of the preceding section, where a lower physiological *H* likewise translated into reduced sleepiness along the identical linear curve, this consistency across pharmacologically and physiologically distinct manipulations provides strong evidence that the threshold distance *H − H*^+^ is a universal predictor of subjective sleepiness, irrespective of the mechanism that modifies its intensity.

Figure 5(c) directly compares the model-predicted and experimentally measured KSS time courses over the 48-h deprivation window. The upper panel shows the placebo condition: both the characteristic monotonic rise during sustained wakefulness and the superimposed circadian modulation are faithfully reproduced by the model. The lower panel shows the caffeine condition, where each 200 mg dose produces a transient dip in KSS that the model captures in both phase and amplitude.

Finally, we use the model to explore how mean sleepiness during prolonged wakefulness depends on caffeine dosage. Figure 5(d) plots the average sleepiness ⟨*H − H*^+^⟩, evaluated after 40 h of total sleep deprivation, as a function of total caffeine intake, for several representative values of the elimination rate constant *k*_*e*_. Despite the well-documented observation that both the elimination rate *k*_*e*_ and the absorption rate *k*_*a*_ decrease slightly with increasing caffeine dose [35, 36], the time-averaged distance ⟨*H − H*^+^⟩ decreases approximately linearly with dose across the range examined. Consequently, the predicted mean KSS also falls linearly with caffeine intake for each fixed *k*_*e*_. A larger *k*_*e*_ (faster clearance) shifts the curve upward, reflecting a weaker cumulative effect for the same total dose; conversely, a smaller *k*_*e*_ (slower clearance) shifts the curve downward, predicting a stronger and more sustained reduction in sleepiness. Our findings suggest that individual differences in caffeine metabolism, captured here by a single pharmacokinetic parameter, can account for much of the inter-subject variability in caffeine efficacy, and that moderate doses in the 100-400 mg range remain within a regime where a simple linear dose-response approximation is adequate.

### Case of Phase axis: circadian modulation of KSS under forced desynchrony

The aging and caffeine analyses of the preceding two sections showed that manipulations of the *intensity* axis–homeostatic parameters (*µ, τ*_*hm*_), steady orexinergic elevation of the threshold margin 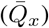, and caffeine-masked drive-shift subjective sleepiness while leaving the linear *H − H*^+^ - KSS mapping intact. Throughout those analyses, however, the circadian pacemaker remained stably entrained to a conventional light-dark cycle, and the threshold *H*^+^ followed its canonical 24-h profile; phase-axis structure was therefore held essentially fixed. A complementary–and arguably more stringent–test of the framework arises when the homeostatic and circadian contributions are experimentally dissociated. In a *forced desynchronization* (FD) protocol, participants are scheduled to a sleep-wake (and light-dark) cycle whose period lies well outside the range of circadian entrainment (e.g. a 20-h or 28-h “day”), so that the endogenous pacemaker free-runs near its intrinsic ~ 24.15-h period while the imposed rest-activity schedule advances at a different rate. As a result, sleep and wake episodes systematically scan across all circadian phases, and the homeostatic drive *H* becomes decoupled from the circadian phase of *H*^+^(*t*). This dissociation is precisely what allows the *phase*-axis contribution of *C* to be isolated: because the “how long one has been awake” component is distributed uniformly with respect to “what time it is,” any residual, phase-locked modulation of sleepiness can be attributed to *H*^+^(*t*) rather than to *H*. If *H − H*^+^ truly constitutes a protocol-invariant sleepiness metric, it should retain predictive validity even when the phase contribution of *H*^+^(*t*) is dynamically decoupled from *H* rather than merely parametrically varied. We therefore asked two questions: (i) does the linear *H − H*^+^ - KSS mapping generalize to FD conditions, in which the circadian pacemaker free-runs and each sleep-wake episode samples a different circadian phase; and (ii) can the threshold geometry cleanly separate the phase-axis contribution of *C*, exposed through the circadian modulation of *H*^+^(*t*), from the intensity-axis effects already demonstrated above?

To address these questions, we used the model to reproduce a canonical forced-desynchrony experiment [19]. After baseline entrainment to a 24 h light-dark cycle (wake 07:00–23:00 under 250 lux; sleep 23:00–07:00 under 40 lux), subjects were placed on a forced-desynchrony (FD) schedule with a non-24 h “day” of period *T* = 20 h. Each FD cycle comprised a scheduled wake episode and a scheduled sleep episode in a fixed 2:1 ratio, i.e. 13.33 h of enforced wakefulness followed by 6.67 h of scheduled sleep. Throughout scheduled wakefulness, the ambient illuminance was held at a constant dim level (*≈* 10 lux) to preclude photic re-entrainment of the circadian pacemaker, whereas scheduled sleep occurred in darkness (0 lux). Because *T* = 20 h lies well outside the entrainment range of the intrinsic circadian period (*τ*_*c*_ *≈* 24.2 h), the imposed sleep-wake cycle and the endogenous oscillator desynchronize and free-run at their respective rates; consequently, successive wake and sleep episodes sample all circadian phases approximately uniformly. The protocol was maintained for *N* FD cycles (here *N* = 30), sufficient for the wake episodes to scan the full 0-360^*°*^ circadian-phase range several times.

Despite this circadian perturbation, the model faithfully reproduced the expected sleep-wake architecture under FD. Figure 6a displays the simulated firing rates of the monoaminergic and VLPO populations, whose flip-flop alternation tracks the FD-imposed *T* = 20 h schedule rather than the endogenous 24 h rhythm. Figure 6b makes this decoupling explicit: the homeostatic process *H* and the circadian process *C* become completely desynchronized, each advancing on its own clock–*H* accumulating and dissipating on the 20 h sleep-wake schedule while *C* free-runs at 24 h–so that the two processes drift continuously through every relative phase. This drift is exactly the condition needed to separate the two axes of the model. Previous sections isolated the *intensity* axis by holding the circadian phase approximately fixed and letting *H* vary; the FD protocol performs the complementary manipulation, scanning *H* uniformly across all circadian phases and thereby exposing the *phase* axis carried by *C*. This separation is made explicit by the threshold itself (Eq. (3)). *H*^+^(*t*) is a purely circadian quantity that translates the instantaneous phase of the pacemaker, through *C*(*t*), into a moving homeostatic set-point. The circadian drive enters Eq. (3) on two routes–linearly through the *−v*_*vc*_*C* term and nonlinearly through the monoaminergic feedback exp[*β*(*v*_*xc*_*C* + …)]–so that *H*^+^(*t*) tracks *C*(*t*) with a phase-dependent gain rather than a constant slope. The threshold distance therefore separates the two axes: intensity and phase, so that, at any fixed level of homeostatic pressure, variation in *H− H*^+^ is driven entirely by the circadian term *H*^+^(*C*(*t*)). Because *H* and *C* run out of phase (Fig. 6b), *H*^+^(*t*) sweeps continuously through its circadian range while *H* evolves on the schedule clock; the sleepiness observed under FD is thus neither a pure intensity effect nor a pure phase effect, but the difference of the two terms which is precisely why *H − H*^+^ remains the appropriate scalar read-out even when the pacemaker is dynamically decoupled.

**Fig 6.**
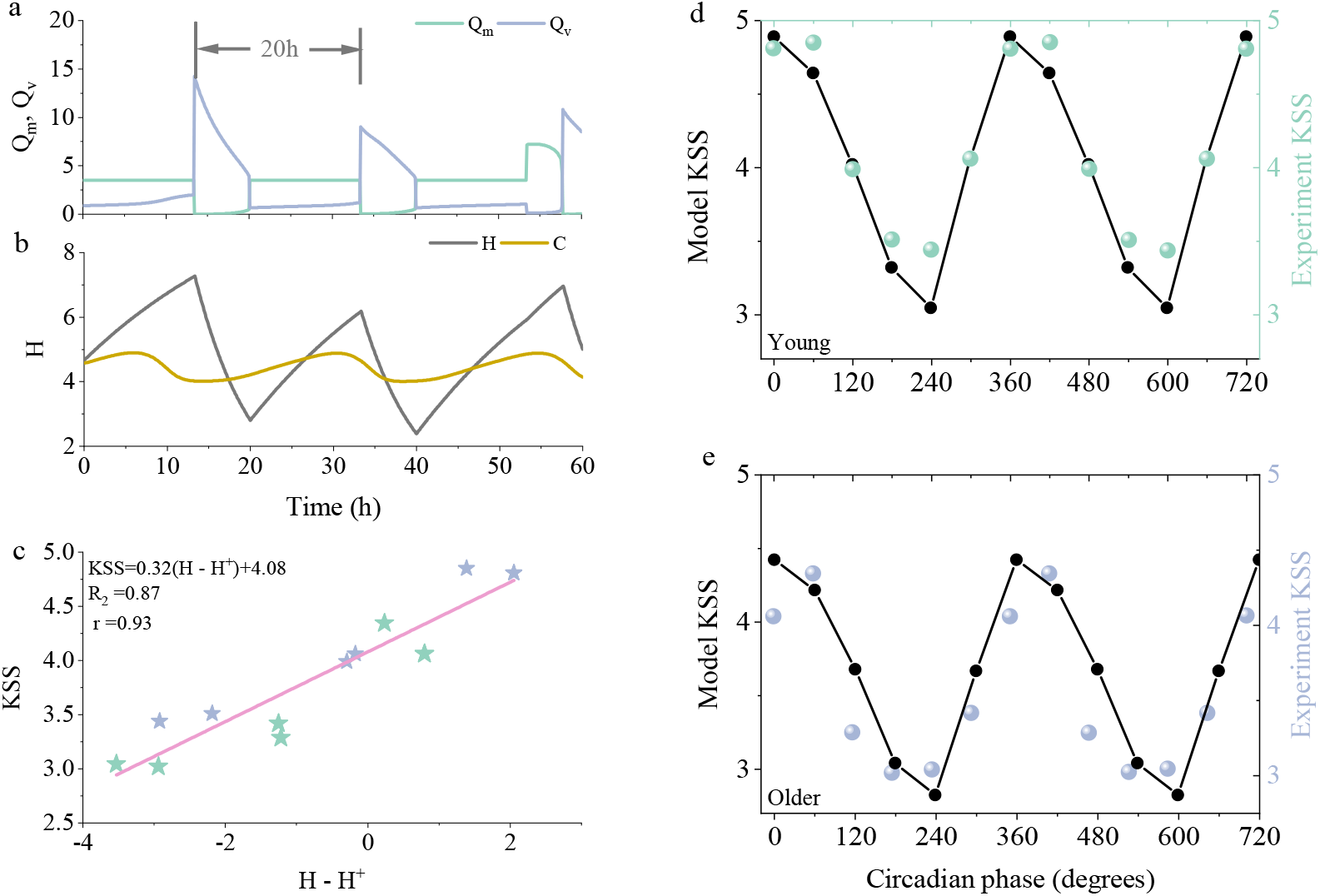
The threshold-distance metric generalizes to forced desynchrony. **(a)** Simulated mean firing rates of the wake-promoting monoaminergic population (*Q*_*m*_) and the sleep-promoting VLPO population (*Q*_*v*_); their alternating dominance tracks the FD-imposed sleep–wake pattern. **(b)** Homeostatic sleep drive *H* (gray) together with the circadian drive *C* (orange); under the *T* = 20 h schedule the two progressively drift out of phase, reflecting desynchronization of the sleep–wake cycle from the free-running pacemaker. **(c)** Subjective sleepiness (KSS) versus threshold distance *H − H*^+^ for older (gray) and young (green) subjects, both collapsing onto a single linear relationship. **(d)–(e)** KSS as a function of circadian phase *ϕ*_CBTmin_ for young (d) and older (e) subjects, respectively.

Critically, pooling data across all FD cycles and circadian phases, the threshold distance *H− H*^+^ maintained a robust linear relationship with subjective sleepiness for both age groups, with the older (gray) and young (green) subjects collapsing onto a single common mapping (Fig. 6c). That one linear law survives a protocol in which the circadian term *H*^+^(*C*(*t*)), not *H*, is the dominant source of variance extends the protocol invariance established in Fig. 2 from the intensity-dominated regime of Figs. 3-4 to a genuinely phase-dominated setting.

Finally, plotting KSS against circadian phase *ϕ*_CBTmin_ (Fig. 6d, young; Fig. 6e, older) isolates the phase term once *H* has been averaged over phase: sleepiness peaks near the circadian trough and troughs near the circadian peak in both groups, tracing out the profile of *−H*^+^(*C*(*t*)) predicted by Eq. (3). The larger phase excursion in the older group reflects their reduced threshold margin, confirming that the age difference reported earlier acts on the same geometric read-out rather than on a separate mechanism.

## Discussions

Building on our previously established linear law KSS = *a* (*H− H*^+^) + *b* [17], here recertified in an orexin-resolved sleep-wake network, we asked what the threshold distance itself resolves into. Across acute deprivation, chronic restriction, aging, caffeine, and FD, the same geometric coordinate continues to track subjective sleepiness; more importantly, its two constituents load onto separable control axes. The sleep-wake network used to compute *H* and *H*^+^ is scaffolding for this geometric claim, not the claim itself.

### Intensity and phase origins of sleepiness

The aging and caffeine studies isolate the *intensity* axis: both act primarily on how large ⟨*H −H*^+^⟩ becomes, while leaving the circadian phase structure of *H*^+^(*t*) largely intact. In aging, reduced *µ* and lengthened *τ*_*hm*_ slow and cap the accumulation of *H*; elevated orexinergic drive 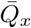 further widens the mean margin by a *steady* upward shift of the sleep-onset boundary–an intensity effect mediated by threshold position, not by circadian phase. Caffeine, by contrast, transiently lowers effective *H* and shifts the operating point down the *same* linear KSS curve. Thus the overall magnitude of subjective sleepiness is set by the time-averaged threshold distance ⟨*H − H*^+^⟩, whether that distance is compressed by moving *H* or by a steady shift of *H*^+^. Consistent with this reading, varying circadian amplitude *r* leaves mean post-deprivation KSS nearly unchanged (Fig. 4a). FD isolates the complementary *phase* axis: when *H* is distributed across circadian phases, residual phase-locked variation in KSS traces *H*^+^(*C*(*t*)) (Figs. 6d,e). Steady threshold shifts (orexin) and circadian threshold dynamics (entrainment/free-run) must therefore not be conflated–both act through *H*^+^, but only the latter constitutes the phase axis. This is precisely the causal statement that fused-drive accounts cannot make: to lower sleepiness intensity one must act on ⟨*H − H*^+^⟩ (sleep, naps, caffeine, or steady wake-holding tone), whereas to reshape its phase one must act on the circadian profile *H*^+^(*t*) (light and schedule regularity).

### Relation to threshold-based sleep modeling

Recent analyses of the two-process framework emphasize that the sleep-onset threshold is not a purely circadian parameter: when related to neuronal mutual-inhibition circuits, mean threshold position reflects network properties and wake-promoting inputs as much as circadian modulation [8, 37]. Our results make that distinction operational for subjective sleepiness. By resolving orexin as an explicit wake-stabilizing population, the mean position of *H*^+^ becomes a physiologically interpretable intensity lever, while the circadian sweep of *H*^+^(*t*) remains the phase lever. The intensity/phase split is therefore not an alternative to the two-process model, but a reading of the threshold geometry that the two-process structure already implies once *H* and *H*^+^ are kept separate and challenged by data [37].

### Implications for personalized sleep-wake management

Threshold-referenced models have already been used to quantify Circadian Sleep Sufficiency (CSS) and to design personalized schedules that reduce daytime sleepiness in shift workers [12, 13]. Those applications show that proximity to a circadian sleep boundary carries information that total sleep time alone cannot provide. What the present dissociation adds is not another sufficiency index, but a criterion for *which lever* to pull. If the dominant problem is an elevated mean sleepiness level–accumulated sleep debt, insufficient recovery between shifts, or a transient need for pharmacological masking–the relevant target is ⟨ *H − H*^+^ ⟩, addressed by extending sleep opportunity, placing restorative naps where the margin is largest, or, when sleep is unavailable, caffeine-like suppression of effective *H*. If instead sleepiness is mainly mis-timed relative to work or social demands–alertness collapsing in the circadian trough during night duty, or residual sleepiness lingering into the commute home–the relevant target is the phase profile of *H*^+^(*t*), addressed by light exposure, schedule regularity, and circadian realignment. In practice, night work and rotating shifts usually mix both deficits: the same roster can raise mean pressure while also placing wake episodes at unfavorable circadian phases [12, 13]. Optimizing only mean alertness, only forced-wakefulness duration during work hours, or only circadian entrainment can therefore pull recommendations in different directions [13]. An intensity/phase decomposition makes those trade-offs explicit before a schedule is chosen, and suggests a concrete next development of existing tools: score candidate sleep-wake and light plans separately for their predicted intensity cost and phase cost, rather than collapsing both into a single alertness scalar. How far that scoring can be individualized–and how it should be fed by wearable sleep and light histories–should be taken up next.

### Calibration and prospective use

The functional form KSS = *a* (*H − H*^+^) + *b* is protocol-invariant, but the fitted slope *a* and intercept *b* vary between cohorts, absorbing inter-individual differences in physiology and KSS reporting. Within a cohort, aging and caffeine effects could be reproduced without re-fitting (*a, b*) (Figs. 3 and 5); across independent studies, recalibration remains necessary. Useful threshold-based predictions likewise require tracking daily changes in sleep pressure and circadian phase–ideally from wearable sleep and light histories–with group-level parameters only as a starting point for individual calibration [12, 13]. A natural next step is therefore subject-specific estimation of a small physiological set–homeostatic rates, orexinergic tone, and circadian amplitude/phase–together with (*a, b*), so that the intensity and phase scores of a proposed schedule can be computed person by person rather than from a population-average model.

### Limitations and extensions

Several limitations qualify these claims. First, KSS is a coarse subjective instrument; whether the same intensity/phase partition holds for objective alertness measures (e.g. psychomotor vigilance) remains to be tested [12]. Second, our circadian module is light-entrained but does not yet exploit richer wearable markers–activity, heart rate, or temperature–that improve phase estimation in irregular schedules [13]; inaccurate phase would corrupt the phase axis even if intensity were well estimated. Third, aging and caffeine were examined as separate intensity probes; stronger tests would orthogonally combine sleep restriction (intensity) with FD or shift schedules (phase) within subjects. Fourth, stimulants enter here only through caffeine’s action on effective *H*; dynamic thresholds under broader stimulant exposure remain an open extension [18, 37]. Finally, disorders in which state-holding capacity itself is altered–most notably orexin-deficient narcolepsy–are natural targets for the same geometric language, but are not modeled here.

In sum, a quantity already used to predict sleepiness and to personalize schedules can be decomposed into intensity and phase origins with distinct physiological controls. That decomposition does not replace existing threshold-based tools; it clarifies how they act, and which lever they should pull, when sleepiness must be managed under the competing demands of sleep debt, circadian phase, aging, and pharmacology.

## Methods

The model architecture is summarized in Fig. 1. Let *V*_*a*_ (*a* = *m, v, x*) denote the mean membrane potential of the MA, VLPO, and Orx populations, respectively, and let *Q*_*a*_ denote the corresponding firing rate. The model is

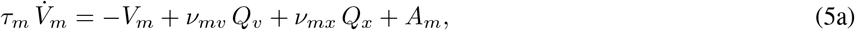

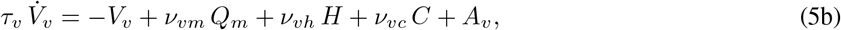

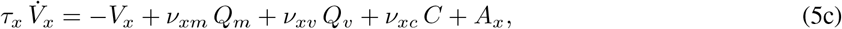

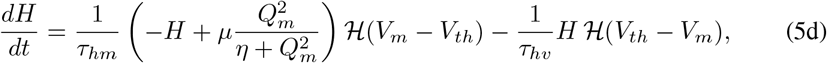

where *τ*_*a*_ are characteristic time constants, *A*_*a*_ are constant background inputs, and *v*_*ab*_ is the effective coupling weight from population *b* to population *a*. Each population firing rate obeys the sigmoidal transfer function

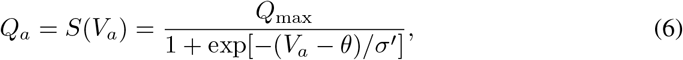

with *Q*_max_ the maximum firing rate, *θ* the mean firing threshold, and *σ*^*′*^ related to the standard deviation *σ* of firing thresholds across the population by 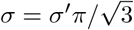 [20].

The homeostatic variable *H* grows as a saturating function of the MA firing rate during wake and decays exponentially during sleep, with the state-dependent switching enforced by the Heaviside function *ℋ* (·) and the threshold potential *V*_*th*_. The coupling *v*_*vh*_ *>* 0 quantifies the excitatory effect of homeostatic pressure on VLPO recruitment [20, 38], while *τ*_*hm*_ and *τ*_*hv*_ govern the buildup and dissipation of *H*, respectively [39]. The constant inputs *A*_*m*_, *A*_*v*_, and *A*_*x*_ aggregate time-averaged drives from sources not modeled explicitly–e.g., cholinergic input to MA, residual excitatory inputs to Orx, and constant offsets of the circadian drive. Nominal parameter values follow the PR model [20], augmented with the additional couplings introduced for the Orx population; all definitions and values are summarized in Table 1.

The signs of the coupling constants encode the architecture defined above: the mutual inhibition between MA and VLPO is enforced by *v*_*mv*_ *<* 0 and *v*_*vm*_ *<* 0; the inhibition of Orx by VLPO and MA is implemented by *v*_*xv*_ *<* 0 and *v*_*xm*_ *<* 0; and the excitatory Orx *→* MA projection is captured by *v*_*mx*_ *>* 0. The dimensionless circadian drive *C* originates from the SCN and is entrained to an effective period of ~ 24.15 h through retinal photic input [40]. It inhibits sleep-promoting neurons (*v*_*vc*_ *<* 0) [41, 42] and excites orexinergic neurons (*v*_*xc*_ *>* 0) [42, 43]. We express *C* in terms of the state variables (*x, y*) of a forced, modified van der Pol oscillator [5],

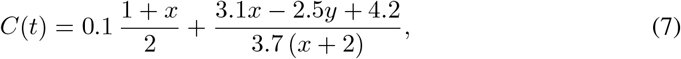

whose coupled dynamics, together with the photic pathway, read

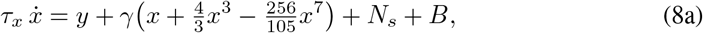

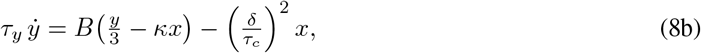

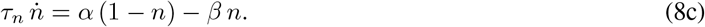

Here *n* ∈ [0, 1] is the fraction of activated photoreceptors. The photic drive to the pacemaker and its associated gain are

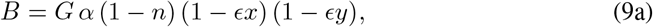

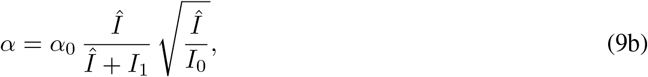

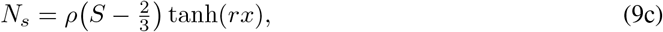

where Î is the effective retinal light input. Both Î and the nonphotic term *N*_*s*_ depend on the behavioral state *S*,

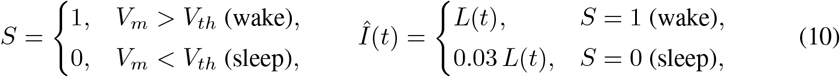

where *L*(*t*) is the ambient illuminance, the factor 0.03 models photic attenuation by closed eyelids, and wake is operationally defined by the threshold crossing *V*_*m*_ *> V*_*th*_. Unless stated otherwise, the circadian parameters take the values 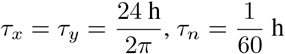, *γ* = 0.13, *ρ* = 0.032, *r* = 10, *G* = 37, *α*_0_ = 0.1, *ϵ* = 0.4, *I*_1_ = 100 lux, *I*_0_ = 9500 lux, *κ* = 0.55, 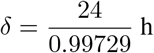, *β* = 0.007, and *V*_*th*_ = *−*2 mV. The light-exposure profile *L*(*t*) is taken from Ref. [5].

In the experiments considered here, the circadian phase is marked either by the phase of the core-body-temperature minimum (CBTmin) or of the plasma-melatonin peak (MELpeak) [5]. To obtain the phase of the circadian pacemaker at time *t* relative to a given marker within circadian cycle *n*, we use

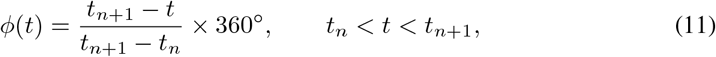

where *t*_*n*_ and *t*_*n*+1_ are the phases of successive circadian markers. These phases are computed following St. Hilaire *et al*. [44], from the phase angle between the two pacemaker variables *x* and *y*: the CBTmin is located where 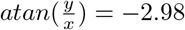 rad, and the marker times are set as *t*_*n*_ = *t*_CBTmin_ + *t*_0_.

Having specified the sleep–wake and circadian machinery, we now introduce caffeine, the intervention of central interest in this work. Caffeine is incorporated via a standard one-compartment pharmacokinetic model [18]. Letting *Z*_*A*_ denote the unabsorbed dose and *Z*_*C*_ the systemic concentration,

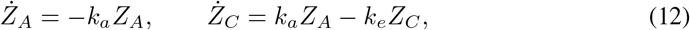

where *k*_*a*_ and *k*_*e*_ are the absorption and elimination rate constants, respectively. A single dose ingested at *t*_0_ with effective amplitude *γ*_*c*_ yields 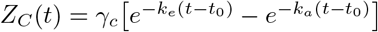 for *t ≥t*_0_, where *γ*_*c*_ absorbs the constant prefactor *k*_*a*_*/*(*k*_*a*_ *− k*_*e*_) of the analytic two-exponential solution. Drug dosages are conventionally expressed in mg/kg, equivalent to parts per million (ppm) by mass; we assume a 75 kg body mass and 130 mg caffeine per cup.

Because caffeine competitively blocks adenosine receptors, it effectively reduces the gain through which homeostatic pressure promotes sleep [45]. To first order in *Z*_*C*_, this is captured by scaling the somnogenic coupling,

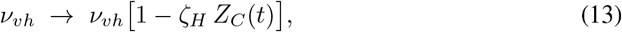

where *ζ*_*H*_ *>* 0 is a constant representing the masking strength. Since the homeostatic term enters Eq. (5) only through the product *v*_*vh*_*H*, this scaling is equivalent to defining an *equivalent* (perceived) homeostatic pressure

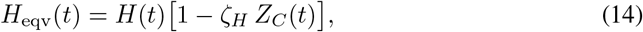

which is the quantity the VLPO effectively senses. Unless stated, otherwise we use *k*_*a*_ = 3.6 h^*−*1^, *k*_*e*_ = 0.16 h^*−*1^, and *ζ*_*H*_ = 0.1.

Because the Karolinska Sleepiness Scale (KSS) is reported at discrete times over an extended protocol, the natural model observable is the *time-averaged* signed distance between the homeostatic sleep pressure and the circadian-modulated sleep-onset threshold,

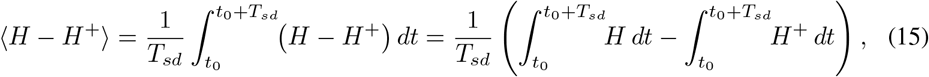

where *t*_0_ is the onset of the deprivation window and *T*_*sd*_ its duration. We evaluate the two integrals in turn.

### Homeostatic component

During sustained wakefulness the homeostatic drive relaxes exponentially toward its waking asymptote *H*_*hm*_,

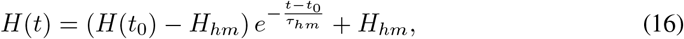

so that its time average over the deprivation window is

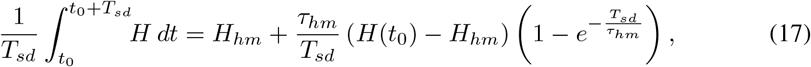

with the waking asymptote *H*_*hm*_ *≈* 0.94 *µ*.

### Circadian-threshold component

Taking the circadian drive to be sinusoidal, *C* = cos(*ω*_*c*_*t*), and expanding the exponential in Eq. (3) to second order in the small quantity *βv*_*xc*_*C*, the mean sleep-onset threshold evaluates to

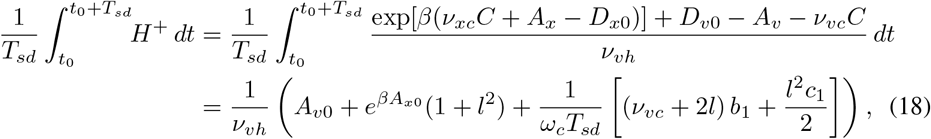

where we have introduced

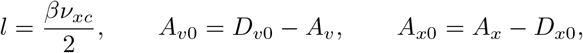

and the boundary terms

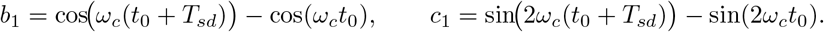

### Combined prediction

Substituting Eqs. (17) and (18) into Eq. (15) yields the analytical approximation

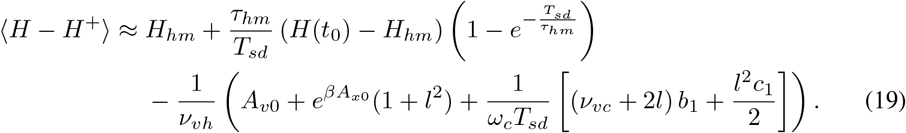

The first line captures the monotonic homeostatic build-up of sleepiness, while the second line encodes the circadian modulation of the threshold. The oscillatory boundary terms *b*_1_ and *c*_1_ enter only through the prefactor 1*/*(*ω*_*c*_*T*_*sd*_) and are therefore *O* (1*/T*_*sd*_): they decay as the deprivation window lengthens and remain subdominant across the physiologically relevant parameter ranges. Consequently, Eq. (19) predicts a well-behaved, frequency-robust average homeostatic load, which we confirm below by direct numerical simulation of the full model.

### Time-averaged equivalent homeostatic pressure with caffeine

We now extend the preceding result to include caffeine, so that the same averaging procedure can be applied to the perceived pressure *H*_eqv_ of Eq. (14). Consider a single scheduled waking episode [*t*_0_, *t*_0_ + *T*_*sd*_] during which one cup of coffee is ingested at time *t*_1_ (*t*_0_ *< t*_1_ *< t*_0_ + *T*_*sd*_). The homeostatic sleep pressure evolves as in Eq. (16), and the systemic caffeine concentration following a bolus of amplitude *γ*_*c*_ at *t*_1_ is

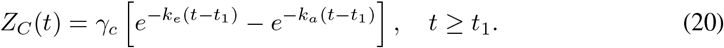

The equivalent (perceived) homeostatic pressure is therefore piecewise,

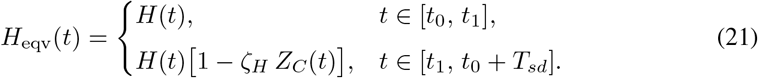

For notational convenience we define

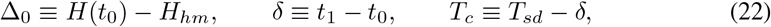

and seek the time-averaged equivalent pressure

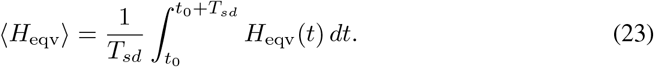

Splitting the domain at *t*_1_, the integral decomposes into a caffeine-free baseline and a caffeine-masking correction,

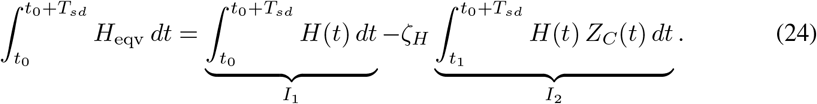

The baseline term reproduces the caffeine-free result,

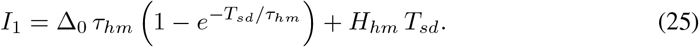

For the correction term, we introduce the local variable *u* = *t −t*_1_, so that *t −t*_0_ = *u* + δ and *u* runs from 0 to *T*_*c*_,

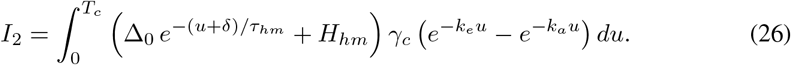

Expanding the product gives

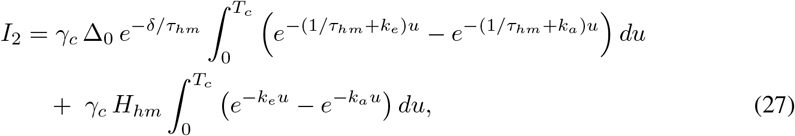

so that, defining 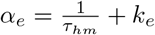 and 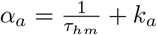,

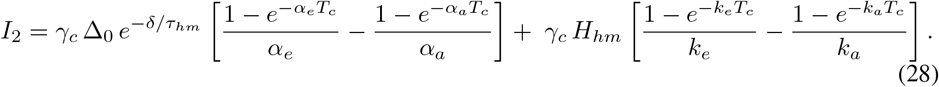

Collecting terms, the time-averaged equivalent pressure reads

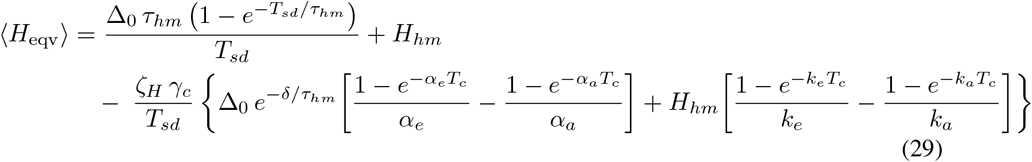

The first line is exactly the caffeine-free average of Eq. (17), while the second, always negative, quantifies the reduction in perceived sleep pressure produced by a single dose. Because this correction is linear in *γ*_*c*_, multiple doses simply superpose, which underlies the approximately linear dose-response relation reported in Fig. 5(d).

## Acknowledgments

This work was supported partially by the Humanities and Social Science Fund of Ministry of Education of China under Grant Nos. 25YJAZH212, the Natural Science Foundation of Zhejiang Province under Grant Nos. LY24A050003, and Jiaxing Public Welfare Project under No. 2025CGZ037.

## Notes

### Competing Interest Statement

The authors have declared no competing interest.

